# A hierarchy in clusters of cephalopod mRNA editing sites

**DOI:** 10.1101/2021.06.16.448723

**Authors:** Mikhail A. Moldovan, Zoe S. Chervontseva, Daria S. Nogina, Mikhail S. Gelfand

## Abstract

RNA editing in the form of substituting adenine to inosine (A-to-I editing) is the most frequent type of RNA editing, observed in many metazoan species. A-to-I editing sites form clusters in most studied species, and editing at clustered sites depends on editing of the adjacent sites. Although functionally important in some specific cases, A-to-I editing in most considered species is rare, the exception being soft-bodied cephalopods (coleoids), where tens of thousands of potentially important A-to-I editing sites have been identified, making coleoids an ideal object for studying of general properties and evolution of A-to-I editing sites. Here, we apply several diverse techniques to demonstrate a strong tendency of coleoid RNA editing sites to cluster along the transcript. We identify three distinct types of editing site clusters, varying in size, and describe RNA structural features and mechanisms likely underlying formation of these clusters. In particular, these observations may resolve the paradox of sequence conservation at large distances around editing sites.

## Background

The mRNA editing process, where an adenine is substituted by inosine (A-to-I editing), is a widespread mechanism of transcriptome diversification in metazoans^1–5^. Inosine is recognized by the cellular machinery as guanine^6–12^, and hence the proteins translated from an edited transcript may be re-coded, thus contributing to the proteome diversity^12–14^. The A-to-I editing is performed by a family of ADAR enzymes, and mutations corrupting ADAR may cause reduction of fitness in model organisms and disease in humans^10,14–18^.

To edit transcripts, ADAR enzymes require specific features of the sequence around editing sites^2,4,5,12,13,19^. Along with the edited adenine itself, a specific, although weak, context is required in nucleotide positions ±1 relative to the edited adenine^13,20–22^. The ADAR enzymes also require edited adenines to be located in RNA helices, which may be parts of complex structures spanning up to over 1kb^2,5^. Thus, the editing at individual sites may be influenced by distant loci, which has been shown directly by the edQTL analysis^23^. However, on average, the span of regions affecting editing at a particular site is about 200–400 nt^13^, as shown by edQTL studies and analysis of conservation levels in regions around editing sites in related species^13,23^. Nonetheless, the context requirements for the A-to-I editing are rather weak, and editing sites have been proposed to form constantly at random points of the genome, especially in structured RNA segments^24^. Adjacently located edited adenines tend to be edited simultaneously^25–29^. Such correlation in human and *Drosophila* has been shown to be mainly due to the involvement of such sites in the same secondary RNA structures^25^. Additionally, editing sites located in coding regions have been shown to be clustered for *Drosophila* and ants, with clusters arbitrarily defined as editing sites located at most at 30–50 nt from each other^30,31^.

A-to-I editing sites are rare in coding regions of most studied genomes, with only minor fractions of them being conserved or functionally important^32–37^. On the contrary, in coleoids (soft-bodied cephalopods, Fig. 1a), not only A-to-I editing is frequent, but conserved sites comprise considerable fractions of all sites, their numbers being orders of magnitude more than in other studied lineages, i.e. mammals and *Drosophila*^13,14,20^. Editing in coleoids, involving up to 1% of all adenines in their transcriptomes, has been suggested to play an important role in proteome diversification, allowing for complex responses to the environment demonstrated by coleoids^13,14^. Along with that, editing sites could have an evolutionary value by rescuing deleterious G-to-A substitutions^38,39^ or by enhancing the heritable trait variance needed for adaptation^22^.

**Figure 1.**
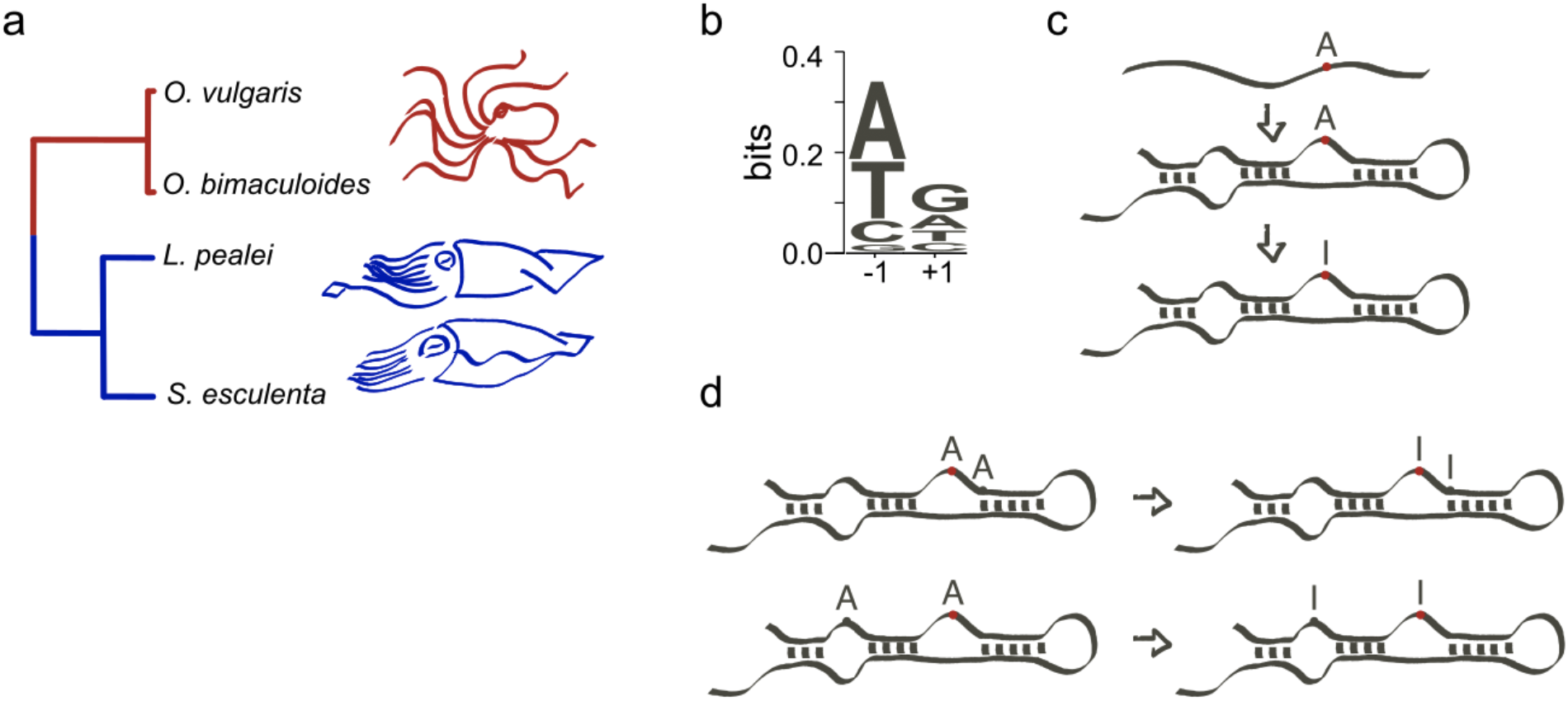
**(a)** Phylogenetic tree of four mollusks (octopuses *Octopus vulgaris* and *O. bimaculoides*, squid *Loligo pealei*, and cuttlefish *Sepia esculenta*) considered in this study. The tree has been taken from TimeTree^54^. **(b)** Sequence context of coleoid A-to-I editing sites. **(c)** ADAR enzymes performing the A-to-I editing require secondary RNA structures. **(d)** The intuition behind this study: closely located adenines should be similar in terms of local RNA structure, and if one of them is edited, the other one also would likely have the necessary prerequisites for the ADAR-mediated editing. Hence, we expect more closely located adenines to be edited simultaneously with a higher probability than more distantly located ones.

By having a large number of A-to-I editing sites, coleoids are a perfect group for studying evolutionary and statistical features of RNA editing. One relevant question is whether coleoid editing sites form clusters and, if they do, what are the average length and structure of the clusters, and what processes underlie cluster formation. As the coleoid editing sites demonstrate same contextual features and secondary RNA structure requirements as mammalian or *Drosophila* editing sites (Fig. 1bc)^13,22,25^, the answer to this question may enhance our understanding of the ADAR action and of the evolutionary and functional mechanisms involved in the emergence of new editing sites.

Here, we rely on four coleoid transcriptomes to show that the levels of association between the A-to-I editing at individual sites strongly depends on the distances between the sites, with the highest correlation observed for immediately adjacent edited adenines (Fig. 1d). We apply multiple, diverse approaches to analyze the distribution of editing sites along transcripts, and identify three distinct types of clusters of coleoid editing sites with sharply different characteristic size ranges. Analyzing local RNA structural features, we observe a tendency of editing sites to be located in putative loops or bulges in secondary RNA structures.

## Results

### Correlated editing

In model species, editing may be correlated if the sites are located sufficiently close to each other^25^. The unusually large numbers of coleoid editing sites allowed us to assess the interplay between co-occurrences of editing states and the distances between editing sites at the single-nucleotide resolution. We used raw RNAseq data (see Materials and Methods) to calculate the correlation of edited states for each pair of editing sites located within the read length interval (~100–150 nt). The correlation coefficient for a pair of edited adenines *E_i_* and *E_j_* given the RNAseq read mapping to transcripts is defined as^25^ (Fig. 2a, Suppl. Figs. S1, S2): 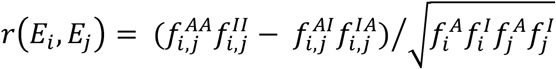, where 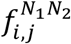 are frequencies of co-occurrences of observed nucleotides *N*_1_ and *N*_2_ (A or I/G) at positions *i* and *j* in the RNAseq read data, and 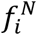 are frequencies of nucleotide *N* in the read mapping data at position *i*. We compared the distributions of *r*(*E_i_*, *E_j_*) for different intersite distances, which we refer to as the *S* values, *S* defined as *j* – *i* (Fig. 2a). The correlations were on average higher for immediately adjacent sites, with mean *r*(*E_i_*, *E_j_*) values further decreasing with the increase of the *S* distance, as in Duan et al., 2017^25^. The power law provided a good fit to the distribution means (*R^2^* = 0.97 to 0.98). The coefficients of the power function inferred from the regression analysis are in the range between −0.28 and −0.22 (Fig. 2a, Suppl. Fig. S1).

**Figure 2.**
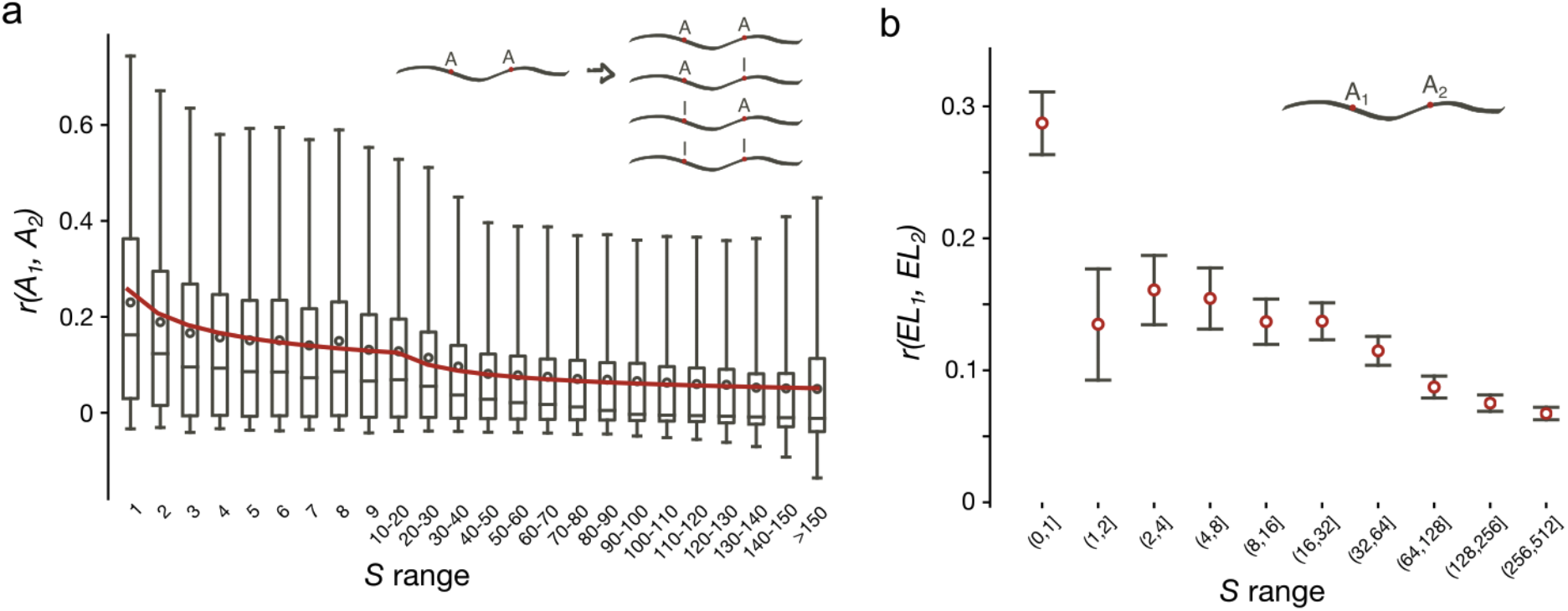
Correlations between various properties of editing sites. **(a)** Distributions of correlation coefficients of *O. vulgaris* editing (*r*) at two sites with respect to the distances between sites (*S*). Boxes represent quartiles, red circles represent the means and the grey lines (whiskers) indicate 95% two-sided confidence intervals of distributions. Red line represents the best power-law fit to the data. **(b)** Dependence of correlations of ELs on the *S* distance, *O. vulgaris* dataset. Red circles mark values of correlation coefficients and grey lines represent Bonferroni corrected 95% two-sided confidence intervals obtained from the *t*-distribution.

The editing level (EL) of an A-to-I editing site is defined as the percentage of transcripts in a sample containing inosine (read as guanine) at the considered site, at the moment of sequencing. As the editing levels of most coleoid editing sites are rather low (<10%), one could speculate that the bulk of associations is lost in the above analyses due to missed low-EL sites that could not be retrieved from the data^21^. Indeed, if we consider sites with EL≥5% (Suppl. Fig. S2a), the average *r*(*E_i_*, *E_j_*) values increase almost twofold, reaching 0.43 for *S*=1. To check whether higher *r*(*E_i_*, *E_j_*) values are not simply a property of efficiently edited sites, we calculated the *r*(*E_i_*, *E_j_*) distributions for sites with the EL≥10% and obtained only slightly larger *r*(*E_i_*, *E_j_*) values, as compared to sites with EL≥5% (Suppl. Fig. S2b). Thus, the association between the A-to-I editing events is indeed strong, especially for adjacently located editing sites.

To check whether the editing state co-occurrence manifests as similarities between ELs, we assessed the correlations between the ELs at individual sites for a series of *S* values (Fig. 2b, Suppl. Fig. S3). For immediately adjacent editing sites (*S*=1), this correlation turned out to be on average twofold larger than for any other *S* (*p*<0.001, the *t*–test). If adjacent sites are not considered, the correlations in ELs do not depend on *S*, being significant (*p*<0.05, the *t*–test) even for quite distantly located sites (*S*>500). The paradox of EL correlations at very large distances may be explained by some transcripts being edited to a higher overall degree than other transcripts. An alternative explanation is as follows. The general variance of ELs in the transcriptome may be decomposed into two summands: the between-transcript variance and the within-transcript variance, the former being the variance of the mean EL values in transcripts, and the latter being the variance of the deviations of ELs from the means in each transcript. If the between-transcript variance is non-zero due to, e.g. low average numbers of editing sites per transcript yielding the estimates of means with high variance, we would observe a baseline correlation for any *S* value, which is simply not defined for sites located in different transcripts.

### Dense editing site clusters (adjacent adenines)

Notably, the correlation between ELs exceeds the baseline only for immediately adjacent editing sites with *S*=1 (Fig. 2b). We consider these sites separately and refer to them as *dense editing site clusters* (DCs) in the general case, and as *paired editing sites* if there are only two adenines per cluster. The observed enhanced positive correlation of editing site co-occurrence for dense clusters (Fig. 2) hints at editing at a focal site being dependent on editing at the immediately adjacent adenine. This could lead to overrepresentation of DCs in the coleoid transcriptomes.

To check whether DCs are indeed overrepresented, we calculated the numbers of sites in DCs separately for each DC size across the coleoid transcriptomes (Fig. 3a, Suppl. Fig. S4). As a control, we constructed a random set of adenines as follows: in each transcript we selected the number of random adenines exactly equal to the number of editing sites in this transcript, and then calculated the site counts in DCs for these random datasets (see Materials and Methods). For all DC sizes, which ranged from two to eight consecutive adenines, the site count in the real datasets was larger than that in the control dataset, the effect being stronger for DCs with larger numbers of adenines (Fig. 3a, Suppl. Fig. S4).

**Figure 3.**
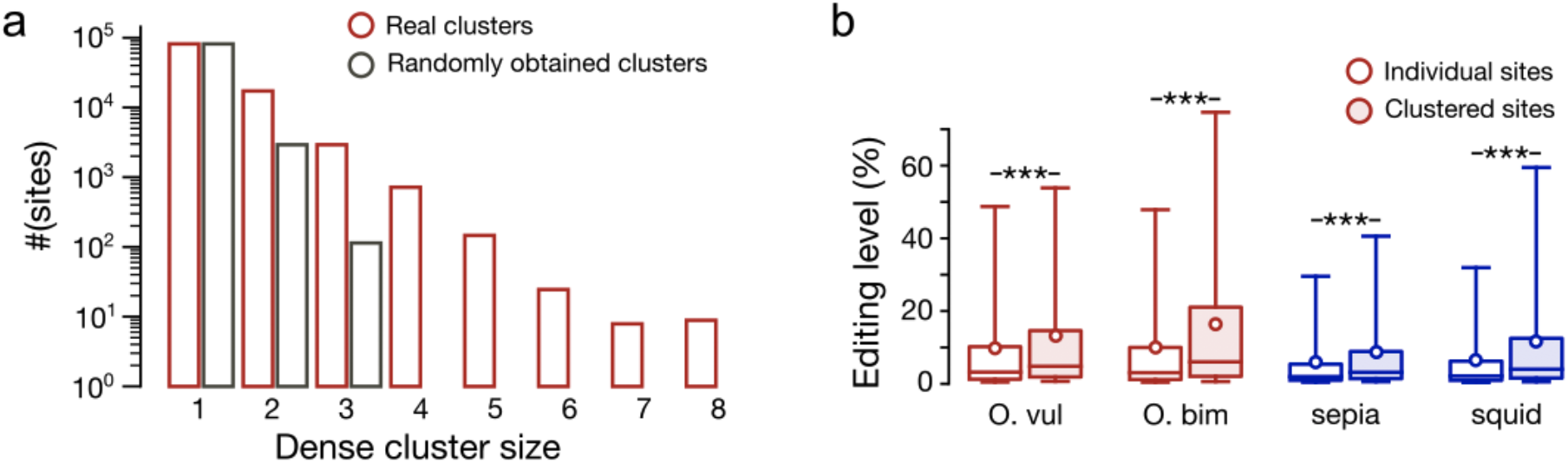
Properties of densely clustered A-to-I editing sites. **(a)** Histogram of dense cluster sizes (nt) for the real *O. vulgaris* editing site dataset (red) and a matching random dataset (grey). **(b)** Comparison of editing levels in densely clustered (filled boxes) sites and in all other sites (white boxes). Three asterisks mark statistical significance of the differences in means (*p* < 0.001, the Mann–Whitney U–test).

Given the observed stronger association of editing at heavily edited adenines compared to that of weakly edited ones (Fig. 2a, Suppl. Fig. S2), one would expect enhanced editing levels of adenines in DCs. Thus, we compared the editing levels in densely clustered sites with those at individual sites (Fig. 3b). The average ELs of sites in DCs were up to 1.6-fold larger than those of individual sites (*p*<7.1 ×10^−158^, the Mann–Whitney U–test) with the fraction of heavily edited sites (EL > 50%) being up to 2.2–fold higher for sites in DCs (*p*<9.6×10^−55^, the χ^2^ contingency test).

### Directionality of dense clusters

As noted above, the strongest association in terms of EL or the co-occurrence of edited states is observed for adjacent editing sites (*S*=1) (Fig. 2ab), with the two-adenine (AA) clusters comprising the vast majority of dense clusters (Fig. 3a). The observed effects may be due to co-operativity of editing, so that, if an adenine is edited, this would enhance the editing context for an adjacent adenine. Moreover, as the editing context is asymmetric (Fig. 1b), we expect probabilities of editing of adenines located immediately up- and downstream from an editing site to differ. To check this hypothesis, we compared the editing levels of upstream and downstream adenines in AA-clusters (Fig. 4a). The ELs at downstream sites were on average 4–6% higher than those of the upstream ones (*p*<1.5×10^−80^, the Wilcoxon signed-rank test) and this result did not depend on the position of the AA-cluster relative to the reading frame of the coding sequence (Suppl. Fig. S5). Thus, our results indicate the AA-clusters directionality.

**Figure 4.**
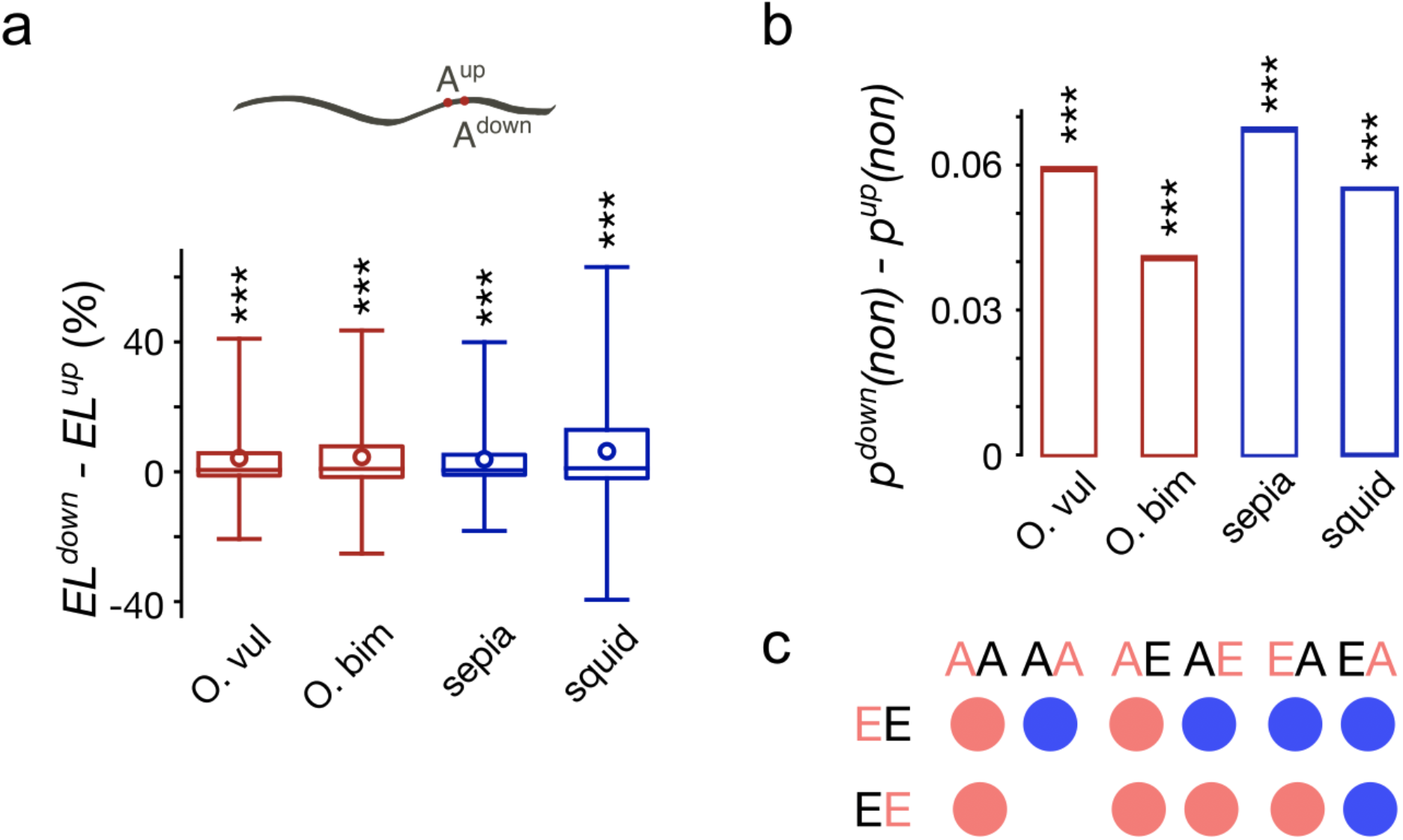
Directionality of dense clusters. **(a)** Distributions of the differences in ELs between down- and upstream editing sites in two-adenine (AA) dense clusters. Three asterisks mark statistical significance of the differences in means (*p*<0.001, the χ^2^ contingency test). **(b)** Differences between the probabilities of down- and upstream editing sites to be non-synonymous. Three asterisks mark statistical significance of the differences in means (*p*<0.001, the binomial test). **(c)** Differences in base-pairing probabilities between paired editing sites (EE) and three types of control AA-dinucleotides (see the text for details). Red color of a letter indicates the nucleotide in a dinucleotide, for which base-pairing probabilities are considered. Red and blue circles show significantly lower and higher base-pairing probabilities for the EE dinucleotide compared to the respective control (the Wilcoxon test *p*<0.05, Bonferroni corrected).

Re-coding (non-synonymous) A-to-I editing in coleoids might be beneficial, as it diversifies the proteome and, consequently, allows for appropriate phenotypic and evolutionary responses to novel environments^13,14,22^. In line with this reasoning, we compared the fraction of re-coding sites among up- and downstream adenines in AA-clusters (Fig. 4b), where both adenines were edited, with the corresponding fractions of re-coding sites in AA dinucleotides, where both adenines were not edited. The probabilities of the downstream sites to be re-coding was higher than those for upstream sites (*p*<3×10^−6^, the binomial test) even accounting for differences in probabilities of editing of adenines in AA dinucleotides.

The differences between ELs and the fractions of re-coding sites of up- and downstream paired edited adenines may be also explained by features of the local secondary RNA structure required for the ADAR action^2^. We assessed the latter explanation by calculating the probabilities of each nucleotide to be involved in secondary RNA structures, which we refer to as base-pairing probabilities (see Methods). For each paired editing site (EE site), we considered the base-pairing probabilities of up- and downstream editing sites separately. As controls, we considered three sets of AA dinucleotides located within ±20nt windows around EE sites: (i) pairs of non-edited adenines (AA sites), (ii) downstream-edited and upstream-unedited adenines (AE sites), and (iii) upstream-edited and downstream-unedited adenines (EA sites). If none of the controls could be obtained for a EE site, it was not considered further (Fig. 4c). As in the case with EE dinucleotides, we considered base-pairing probabilities in control dinucleotides separately for up- and downstream nucleotides.

Firstly, we observed the base-pairing probabilities of downstream adenines in EE sites to be significantly higher than those of upstream adenines (Wilcoxon p < 2.6×10^−39^). The dependency of base-pairing probabilities on the adenine position in a dinucleotide extends to the comparison of base-pairing probabilities of EE dinucleotides with those of control dinucleotides (Fig. 4c, Suppl. Tab. S1), where the downstream adenine seems to be generally less structured than the upstream adenine. Additionally, positions of editing sites in the control sets largely and consistently affect the results: AE controls are generally more structured than the EA controls (Fig. 4c). Thus, the downstream adenines in EE clusters are edited more frequently, are more likely to be re-coding if edited, and are less likely to be involved in secondary RNA structure (Fig. 4).

### Medium-range clusters of editing sites

Correlations in the editing state co-occurrence for *S* values larger than 1 (Fig. 2a) hint that A-to-I editing sites may cluster not only in the form of DCs. Thus, we checked how the distance to the nearest editing site affects the probability of adenine editing (Fig. 5a). We introduce the measure *S*\* defined as the distance between two edited adenines such that no other edited adenine is located between them, and consider the deviation of the observed *S*\* distribution from the expected one (Fig. 5a). The expected distributions were calculated on randomly generated datasets described above. For all considered coleoid species, the observed and expected *S*\* distributions differ significantly only for windows of up to 18 nucleotides (*p*<0.01, the χ^2^ test with the Bonferroni correction), thus suggesting a direct dependence of editing events within the 18nt distance.

**Figure 5.**
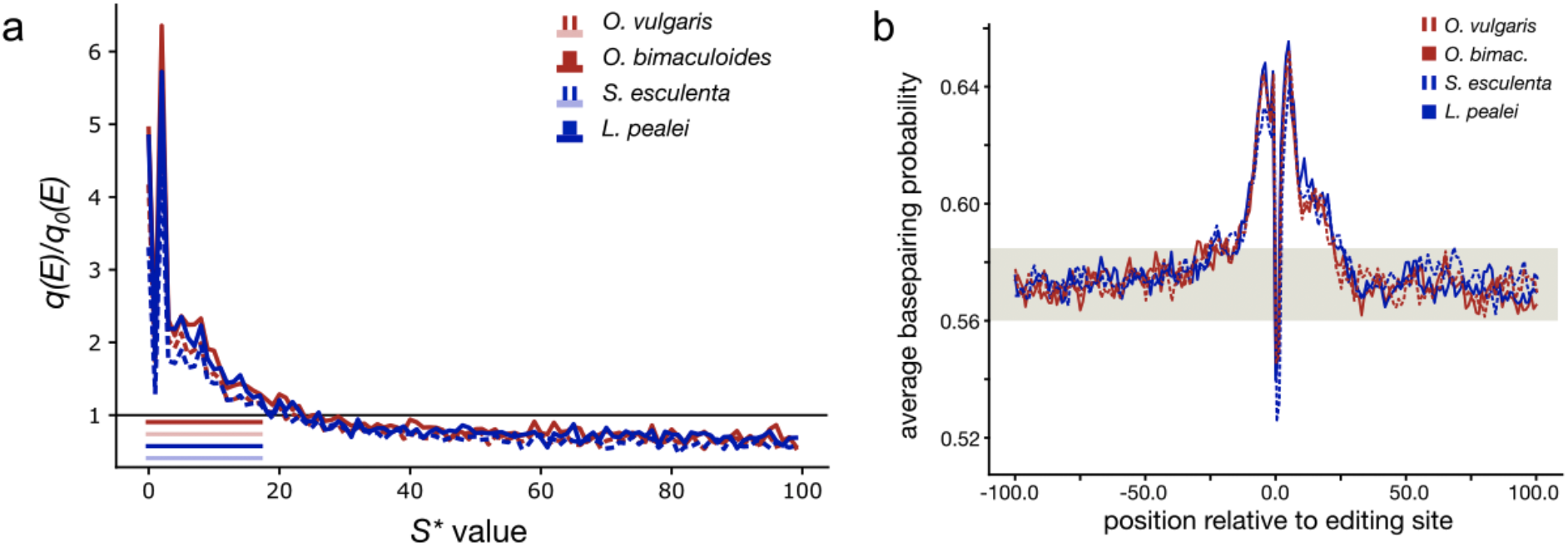
Properties of medium-range clusters of editing sites. **(a)** Deviation of the editing probabilities of adenines located near editing sites (*p*(*E*)) from the respective expected probabilities (*q*_0_(*E*)) as dependent on the *S*\* values. The colored stripes in the lower left corner represent the *S*\* value ranges on which *q*(*E*) are significantly higher than *q*_0_(*E*) (*p* < 0.01, the χ^2^ contingency test, Bonferroni corrected) **(b)** Average base-pairing probabilities in the regions centered at editing sites in four coleoid species. The gray stripe marks the base pairing probability range in regions distant from editing sites (>200 nt), considered as noise. The values above the noise (the central peak) describe the putative average RNA structure around editing sites; the width of the peak is the average size of the structure. The dip in the middle is caused by generally low base-pairing probabilities of edited adenines.

As noted above, A-to-I editing requires secondary RNA structures to be formed around the edited adenine^2,10,22^. Hence, the observed clustering of editing sites may be explained by common RNA structures at clustered sites. Thus, we have assessed the average size of a local secondary RNA structure by analyzing average base pairing probabilities of nucleotides around editing sites (Fig. 5b). The average RNA structure size for each coleoid species is determined as the average width of peak in pairing probabilities of nucleotides centered at editing sites; the peak is defined at the region where the average base-pairing probabilities are greater than those of nucleotides distant from editing sites. So defined peaks for all four considered coleoid species fall in the range 32–45 nt, which is consistent with the above estimate of the distance at which an edited adenine influences the probability of editing of a neighboring adenine, which is 2×18 nt = 36 nt (Fig. 5a). Thus, the correlated editing of adenines located sufficiently close to each other indeed may be caused by common local secondary RNA structures. Moreover, as there is a higher probability of editing of adenines located in the vicinity of editing sites, editing sites should cluster along the transcript, forming what we call medium-range editing site clusters.

The previously obtained result about a subset of heavily edited and re-coding sites being less likely involved in secondary RNA structures (Fig. 4) seemingly contradicts earlier observations that these sites tend to reside within structured regions^22^. This controversy was resolved by nucleotide-resolution structural analysis of regions around editing sites. For each edited adenine we sampled the nearest non-edited adenine as a control and assessed the site and control base-pairing probabilities (Suppl. Fig. S6). The base-pairing probability of control sites turned out to be larger than that of editing sites, the effect being stronger for sites with large ELs (Suppl. Fig. S6a). Moreover, the energy of the local secondary RNA structure was lower for editing sites compared to that of control ones (Suppl. Fig. S6b), confirming that the RNA structure around editing sites is more stable than that at the editing sites themselves. The observed pattern suggests that editing sites tend to reside in loops or bulges, i.e. in non-paired regions surrounded by stable helices.

### Long-range clusters of editing sites

Earlier studies of coleoid editing sites demonstrated relatively higher sequence conservation in intervals of ±100–200 nt relative to conserved editing sites^13^ and a correlation between differences in the editing levels at homologous sites and the number of mismatches in the ±100 nt region^22^. These two consistent estimates indicate that editing at focal sites depends on ±100–200nt context, which exceeds the size of medium-range cluster sizes, established above as of 32–45 nt (Fig. 5).

Medium-range clusters have been identified by probability measures. A complementary approach is the comparison of real and expected *S* values, *S* being the distance (in nucleotides) between two edited adenines located in a single transcript, regardless of other possible editing sites between them. As with dense and medium-range clusters, the null model for *S* values was derived from a random set of adenines with the per-transcript number of editing sites preserved (see Materials and Methods). We have observed that the distribution of distances, *S*, calculated for known coleoid editing site sets is bimodal with a high and distinct peak at 1, reflecting overrepresentation of edited adenines in dense clusters (Fig. 6, red curve). Having calculated distances *S* using the randomized set of adenines, we have observed strong and highly significant differences between the real and control *S* distributions (*p* < 2.2×10^−308^, the Kolmogorov–Smirnov test, Fig. 6). At that, the differences are limited to distances *S* smaller than approx. 100–200 nt (Fig. 5, Suppl. Fig. S7), consistent with the earlier observations mentioned above^13,22^, and yields long-range editing site clusters at the scale of 200–400 nt.

**Figure 6.**
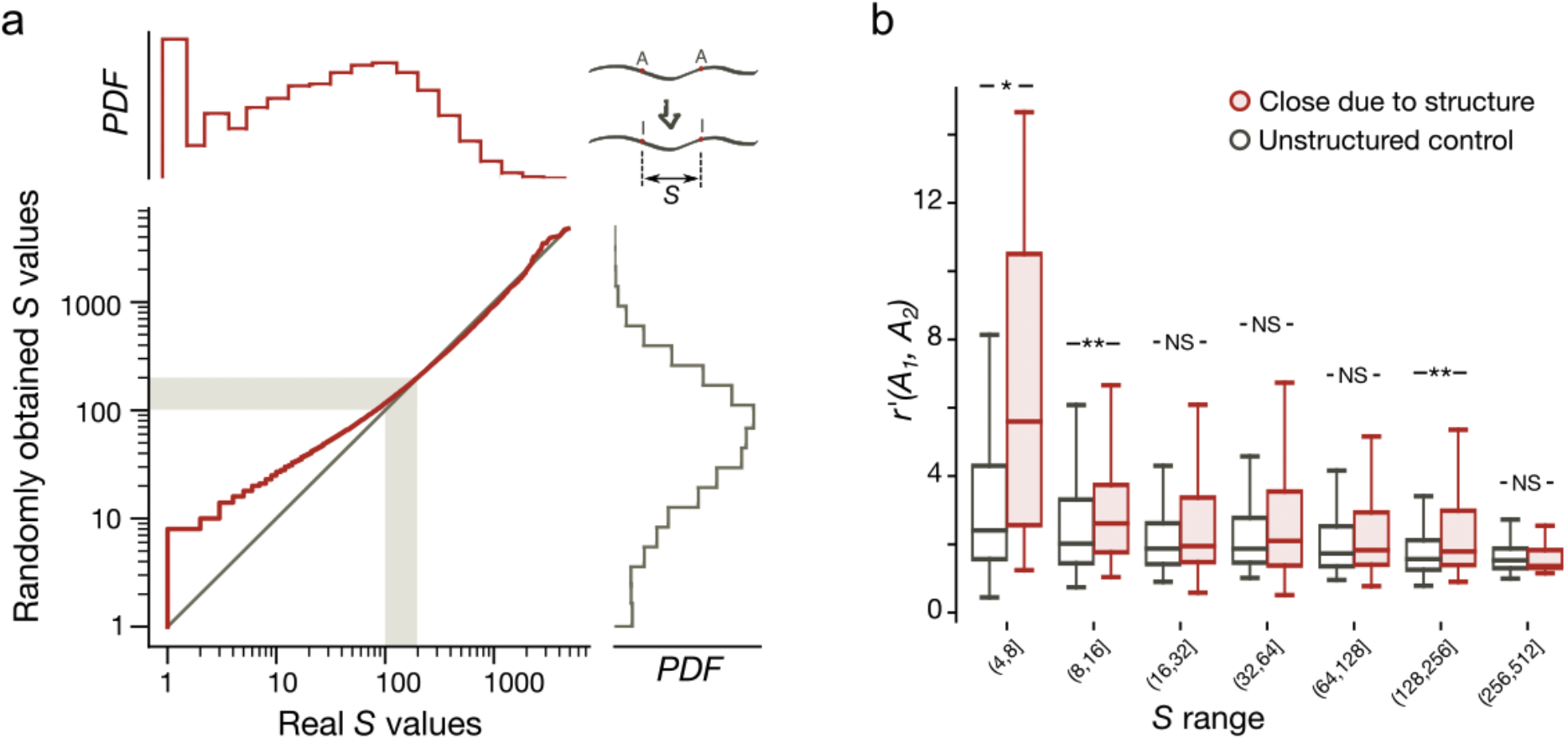
Long-range editing site clusters. **(a)** Distribution of *S* distances for *O. vulgaris*. The real editing site set (red histogram) *vs*. randomly selected adenines (grey histogram), see the text for details. The red line is the plot of dependence between the real and the randomly obtained *S* values in arrays sorted by the distance *S*. The grey diagonal represents the expected dependence form *y=x*. Grey stripes represent the boundary of the possible span of regions affecting editing sites^13,22^. **(b)** Distributions of the *r*′(*A_i_*, *A_j_*) values calculated for the structurally close editing sites (red boxes) and for the control site pairs with no predicted secondary RNA structure between the sites in a pair (grey boxes) (see the text for details). Asterisks mark statistical significance of differences of means calculated using the Mann-Whitney U-test with the Bonferroni correction for binning. Two asterisks indicate *p*<0.01; one asterisk, *p*<0.05, NS, non significant.

To understand the mechanisms yielding long-range clusters, we applied a relaxed definition of RNA structure spanning over a pair of edited adenines. We considered pairs of adenines brought close to each other in space by secondary RNA structure (see Methods). As a control, we considered pairs of sites such that no secondary RNA structure could be identified between them (Fig. 6b). As a measure of co-operativity of editing, we employed the formula: 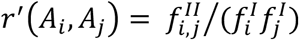, where 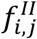 is the frequency of co-editing at a pair of sites *i* and *j*, and 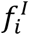 and 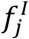 are the individual frequencies of editing at the respective sites. Editing sites brought close by secondary RNA structures were generally more co-operative (*p*=7.8×10^−7^, the Mann-Whitney U-test) than the control sites, with the sites at distances 4–16 nt and 128–256 nt exhibiting significant increase in co-operativity when brought close by secondary RNA structure (*p*<0.05, the Mann-Whitney U-test with the Bonferroni correction for binning) (Fig. 6b). This result indicates the effects on co-operativity at characteristic long-range cluster sizes to be brought about by secondary RNA structures. These structures could be expected to be rather weak on average, as the structural potentials of nucleotides at distances from editing sites larger than 36 are indistinguishable from noise (Fig. 5b).

## Discussion

### Cooperativity of RNA editing

A-to-I editing sites in coleoid genomes tend to cluster. The strength of correlations in the editing state co-occurrence clearly depends on the distance between the sites, well described by a power law. One explanation is provided by the common secondary RNA structure formed around closely located editing sites. However, the common RNA structures do not explain the inosine co-occurrence observed in our (Fig. 2a) and in other^25^ studies. Indeed, suppose an adenine is edited due to the local RNA structural features. The local structure would generally enhance the probability of editing of adjacent adenines^24^, however, editing at an adjacent site would not depend on the editing at the considered site, unless the RNA structure has changed due to the first act of editing. Thus, no correlations would be observed. This prompts for a dynamic explanation based on changes of editing probabilities near the focal site introduced by editing at that site. We consider the following two scenarios: (i) ADAR enzymatic action at adjacent editing sites is co-operative, manifesting as simultaneous adenine editing dependent on the linear distance between the edited adenines and (ii) inosine produced by editing at one site stabilizes the existing local secondary RNA structure or even causes RNA to fold in a different manner, hence enhancing the probabilities of editing at nearby adenines.

The former explanation presumes that ADAR enzymes can edit multiple sites in a series of enzymatic acts, this ability being dependent on intersite distances. This is indirectly supported by the fact that different ADAR subunits show enzymatic cooperativity for substrate binding^40^. Similar effects are observed e.g. in the case of co-operative phosphorylation of adjacent amino acids in proteins, where clusters of phosphorylated residues form due to the enzymatic features of phosphatases^41–43^. That, however, does not explain the prevailing editing state co-occurrence in the adjacent adenines, as two ADAR subunits may not physically edit two consecutive adenines simultaneously^44^. However, the latter effect can be explained by slippage of the ADAR RNA-binding domain on the RNA sequence, resulting in editing of the adjacent adenine.

In the RNA-centered model, the seeming co-operativity of A-to-I editing of adjacent sites is attributed to the reinforcement of the local secondary RNA structure, which would increase the probabilities of editing at adjacent or closely positioned adenines. Inosines form base-pairs with cytosines, the I-C base pair being isosteric to, but slightly less stable than the G-C pair^45^. Together with our observation about edited adenines being on average less structured than control, nonedited adenines close to editing sites (Fig. 4ab), this points to a possibility that editing at a focal site changes the local RNA structure pattern, reinforcing the local propensity towards secondary structure, and hence promotes editing at adenines in the vicinity. We could not test this explanation computationally due to insufficient data on structural features of inosines^45^.

### Structure of dense clusters

Downstream adenines at EE-clusters are more heavily edited and more frequently non-synonymous (Fig. 3cd). One possible explanation for this comes from the editing site sequence context (Fig. 1b), where adenine is preferred upstream of editing sites, and, as the two adenines in DCs possess common local secondary RNA structures, the downstream site should be edited more frequently. However, this does not explain these adenines being recoding more often; the latter observation asks for both more editing data from additional species for signs of selection and for experiments on functional significance of these editing events.

### The range of influence of editing sites

Previous studies have established the linear lengths of RNA structures associated with A-to-I mRNA editing to be of various sizes ranging from rather short structures^19^ to complex formations spanning over large fragments of the transcript^2^. In coleoids, conserved regions around conserved editing sites span on average 100– 200 nt in each direction^13^. Accordingly, clustering of edited adenines obtained from the *S* value analysis and the analysis of structurally close edited adenines is observed at up 100–200 nt and up to 256 nt, respectively (Fig. 2a). However, the analysis of adenine editing probabilities in the vicinity of edited sites (Suppl. Fig. S2) and the analysis of base-pairing probabilities in the regions around edited adenines (Fig. 4c) have yielded different and consistent estimates of 36 nt and 32– 45 nt, respectively. This indicates a hierarchy in the cluster structure, with relatively large, diffuse clusters spanning up to ~400 nt yielded possibly by weak secondary RNA structures associated with editing sites, which span up to 256 nt (Fig. 6). Smaller, however more stable structures spanning up to 45 nt yield the intermediate level of clustering (Fig. 5). Finally, the local features of RNA structure, e.g. loops and bulges, confer the strongest association in terms of editing, which manifests as clusters of adjacent edited adenines (Fig. 2, Fig. 4c).

## Materials and Methods

### Data

In the present study, we employed the previously published transcriptomes^13^ of *O. vulgaris, O. bimaculoides, S. esculenta*, and *L. pealei* along with the publicly available coleoid editing sites data^13^. The corresponding transcriptomic read data summarized in Suppl. Tab. S2 were downloaded from the SRA database.

### Calculation of *S* values

*S* values were calculated as nucleotide distances between edited adenines on transcripts. Along with *S* values calculated for actual editing sites, we calculated *S* values for randomly selected adenines. To eliminate the biases caused by factors such as the higher accuracy of editing sites prediction in highly expressed transcripts or the general tendency of some transcripts to be edited more frequently than others, we have randomly selected in each transcript the number of adenines equal to the number of editing sites it contains. The *S*\* values were calculated as nucleotide distances between subsequent edited adenines, i.e. for pairs of editing sites with no edited adenines between them.

### Editing state co-occurence

To infer the tendency of edited states to co-occur in transcripts, we have calculated the Pearson correlation^46^ of edited state occurrences for all pairs of edited adenines located within the window with the radius equal to the read length.

The reads were mapped onto the transcripts with the bowtie2 package^47^ using the – sensitive-local alignment mode. We filtered out read alignments that did not contain regions of continuous read mappings larger than half the read length. The resulting alignment files were further processed with a set of ad hoc scripts and the numbers of editing state occurrences for each considered site pair were calculated.

### RNA structural annotations

To estimate the propensity of sequences to form RNA secondary structure, we have calculated the structural potential Z-scores for each nucleotide using the RNASurface program^48^. *Z-score* is defined as *Z* = (*E* – μ)/σ with *E*, μ and σ being the minimal free energy of a considered cequence, mean and standard deviation of the free energy distribution of shuffled sequences with preserved length and average dinucleotide composition, respectively. RNASurface was run with the maximal and minimal sliding window length set to 350 and 20, respectively. For each position, *Z-score* was inferred as the minimal *Z-score* of all windows containing it.

The base pairing probabilities were calculated with the plfold algorithm of the Vienna package^49^ with –W and –L parameters set to sequence lengths and –cutoff parameter set to 0.0.

For the analysis of editing sites brought close by secondary RNA structures, all possible pairs of editing sites for each transcript were considered. For Fig. 6b, every such pair was assigned to one of the three groups: “close due to structure”, “distant, unstructured”, or “intermediate”. Two editing sites were considered close due to the structure if the distance between them in the structure was less than the distance between them taken by sequence, and, additionally, the distance in structure was less than eight nucleotides. The pair of sites were considered distant if the distance between the sites in the structure was equal to the distance by the sequence or was more than 40 nucleotides. The distance in the structure for the pair of sites was computed as the minimal distance between them in the graph of the transcript with all the potentially paired base pairs and all nucleotides adjacent in the sequence connected by edges. The graph was obtained using the Vienna RNAplfold program with the 0.8 cut-off for the pairing probability^49^.

### Statistics

The distributions of *S* values were compared using the two-sample Kolmogorov-Smirnov test^50^. Correlations between the editing levels at pairs of sites were assessed with the Pearson correlation coefficient; the confidence intervals and the significance of each correlation coefficient were inferred using the t-test with the Bonferroni correction^51^ for multiple testing. Editing levels, the distributions of correlation coefficients, and the distributions of structural potential Z-scores were compared with the Mann-Whitney U test^52^. Probabilities of upstream and downstream clustered editing sites to be synonymous as well as probabilities of editing sites to be located in specified codons or in protein disordered regions were compared with the binomial and the χ^2^ contingency tests. The editing levels at upstream and downstream editing sites were compared with the Wilcoxon signed-rank test^53^.

The grouping of *S* values with respect to the differences in correlations between edited states on transcripts was performed using the Mann-Whitney U test: for each pair of correlation arrays corresponding to different *S* value ranges, the Mann-Whitney statistic was calculated, and groups of *S* value ranges were further defined as the groups of sequential ranges differing insignificantly from each other.

### Code availability

All data analyses were performed in Python 3.7. Scripts and data analysis protocols are available online at https://github.com/mikemoldovan/coleoidRNAediting2.

## Supporting information

File containing all supplementary data

