## Supplementary material for "A hierarchy in clusters of cephalopod mRNA editing sites": File containing all supplementary data

### Supplementary materials for
### “Clusters of mRNA ADAR editing sites in soft-bodied cephalopods”

#### Supplementary Figures

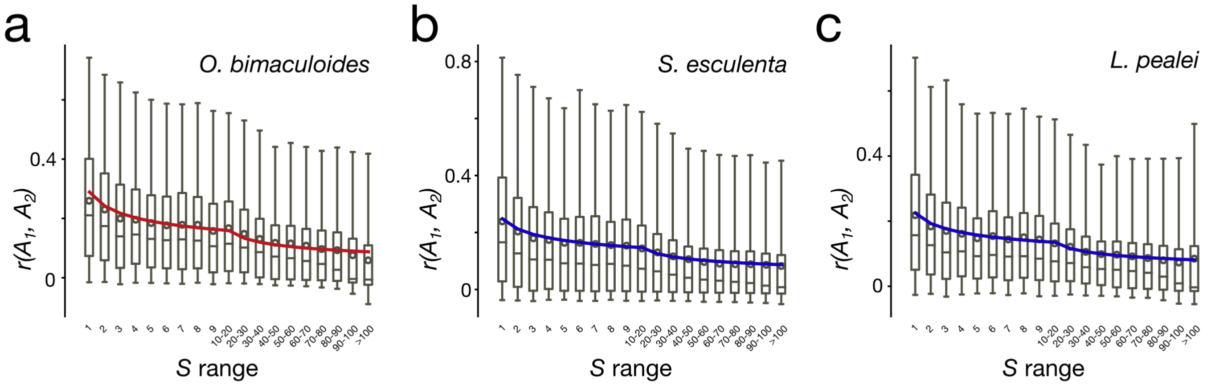


**Supplementary Figure S1 | The distributions of correlation coefficients of coleoid editing at two sites with respect to the distances between sites.** Boxes represent the quartile borders, red circles represent the means and the grey lines indicate the 95% two-sided confidence intervals of the distributions. Red and blue lines represent the best power fits to the portrayed data. **(a)** *O. bimaculoides* **(b)** *S. esculenta* **(c)** *L. pealei*. Notation as in Fig. 2a.

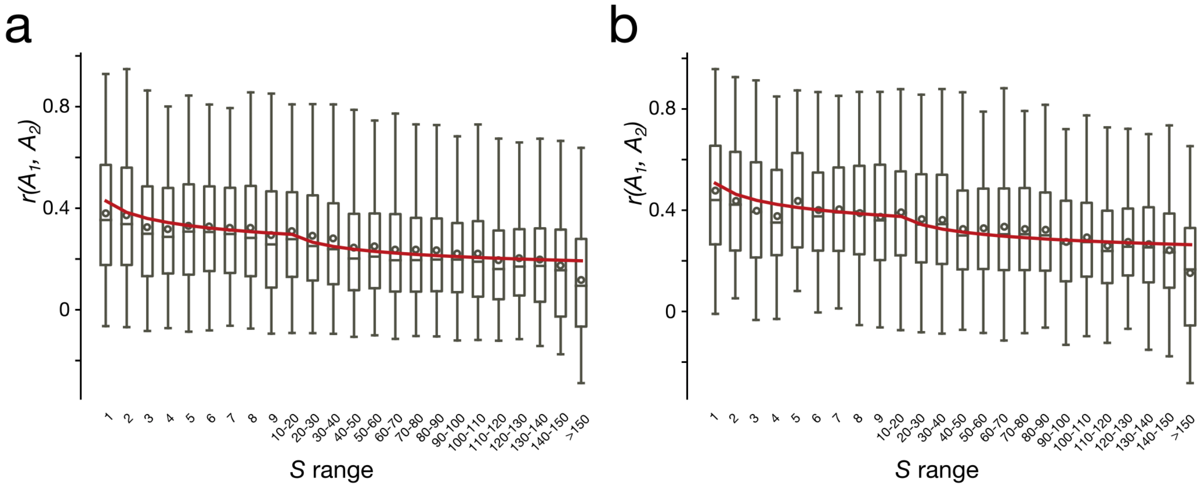


**Supplementary Figure S2 | The distributions of correlation coefficients of *O. bimaculoides* editing at two sites with respect to the distances between sites for different minimal editing level threshold values.** Boxes represent the quartile borders, red circles represent the means and the grey lines indicate the 95% two-sided confidence intervals of the distributions. Red and blue lines represent the best power fits to the portrayed data. **(a)** Threshold set to 5% **(b)** Threshold set to 10%. Notation as in Fig. 2a, Suppl. Fig. S1.

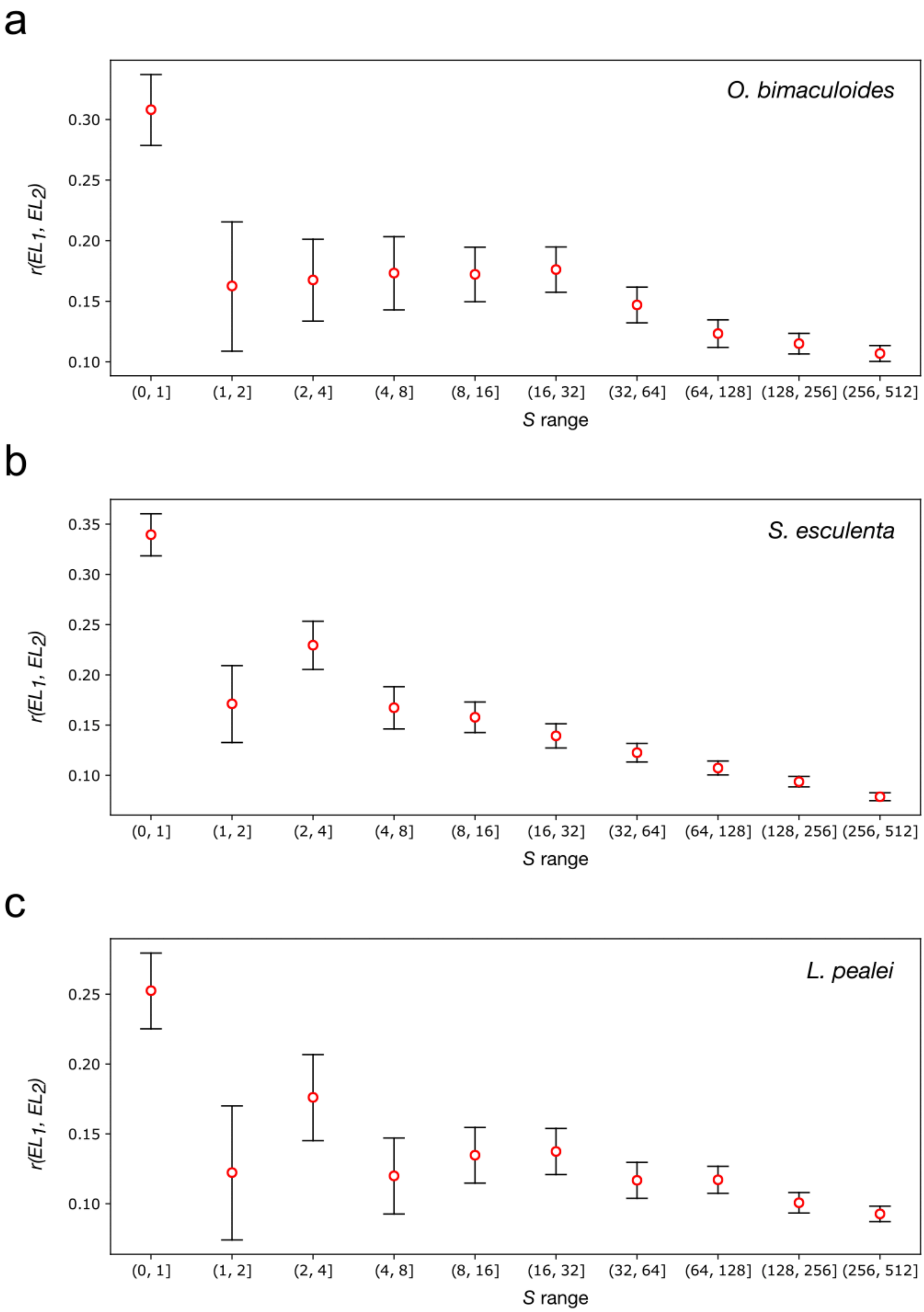


**Supplementary Figure S3 | The dependence of the correlations of ELs on the *S* values for the considered coleoid A-to-I editing site datasets**. The red circles mark the values of the correlation coefficients and the grey lines represent the Bonferroni corrected 95% two-sided confidence intervals obtained from the *t*-distribution. **(a)** *O. bimaculoides* **(b)** *S. esculenta* **(c)** *L. pealei*. Notation as in Fig. 2b.

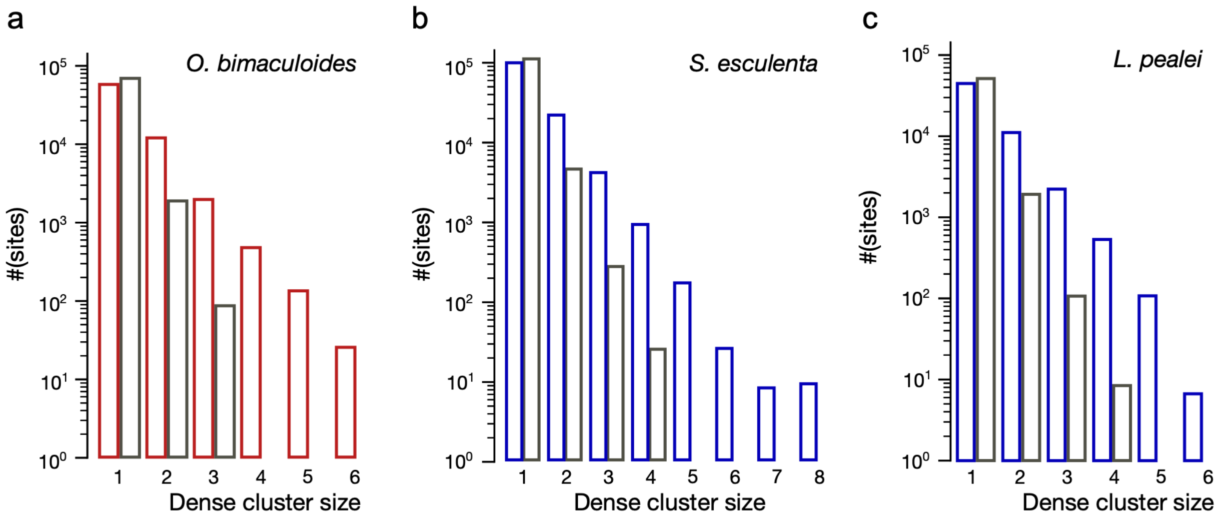


**Supplementary Figure S4 | Histograms of dense cluster sizes (nt) for the real coleoid editing site datasets (red and blue) and the corresponding randomly obtained ones (grey). (a)** *O. bimaculoides* **(b)** *S. esculenta* **(c)** *L. pealei*. Notation as in Fig. 3a.

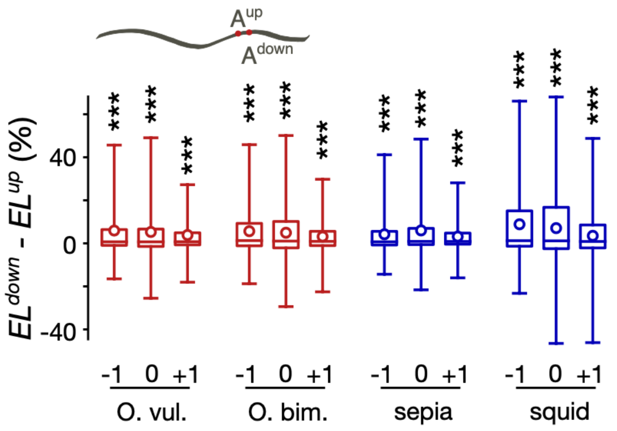


**Supplementary Figure S5 | The distributions of the differences in ELs between down- and upstream editing site in two-adenine dense clusters obtained for the three reading frames.** Three asterisks mark statistical significance of the differences in means ($p < 0.001$, Chi-squared contingency test). The values -1, 0 and +1 below indicate the coordinate of the dense cluster reading frame relative to the predicted protein-coding frames in coleoid transcriptomes.

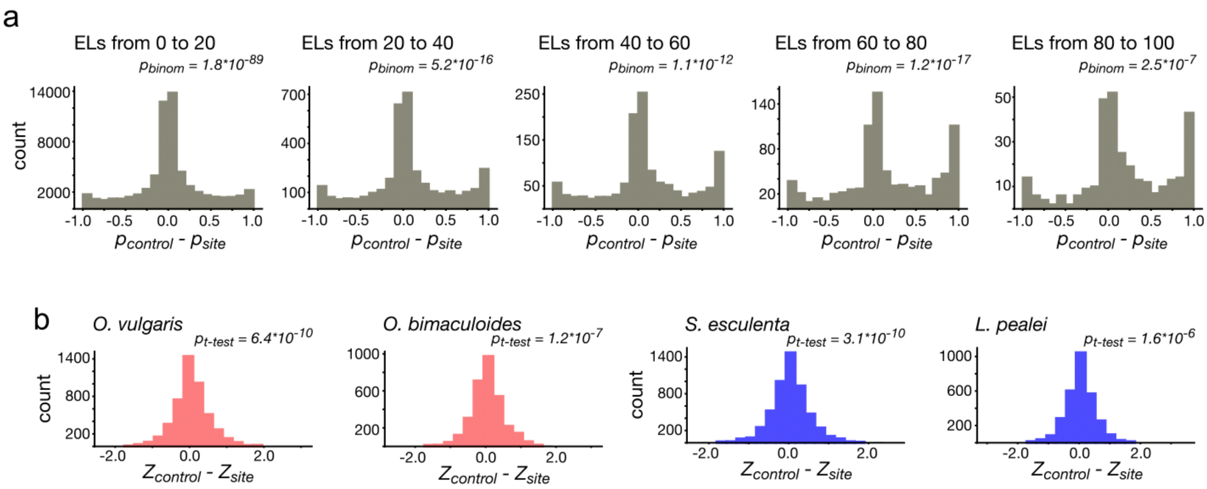


**Supplementary Figure S6 S8 | Structural properties of coleoid A-to-I editing sites. (a)** Differences in base pairing probabilities between A-to-I editing sites and nearest unedited adenines used as controls for different editing level (EL) intervals in *O. vulgaris*. The significance (*p*-values, the binomial test) of the site–control differences is shown. **(b)** Differences in RNA secondary structure free energy between editing sites and control adenines in four coleoid species. The significance (*p*-values, the *t*-test) of the control sites structural potentials being larger than those of editing sites is shown.

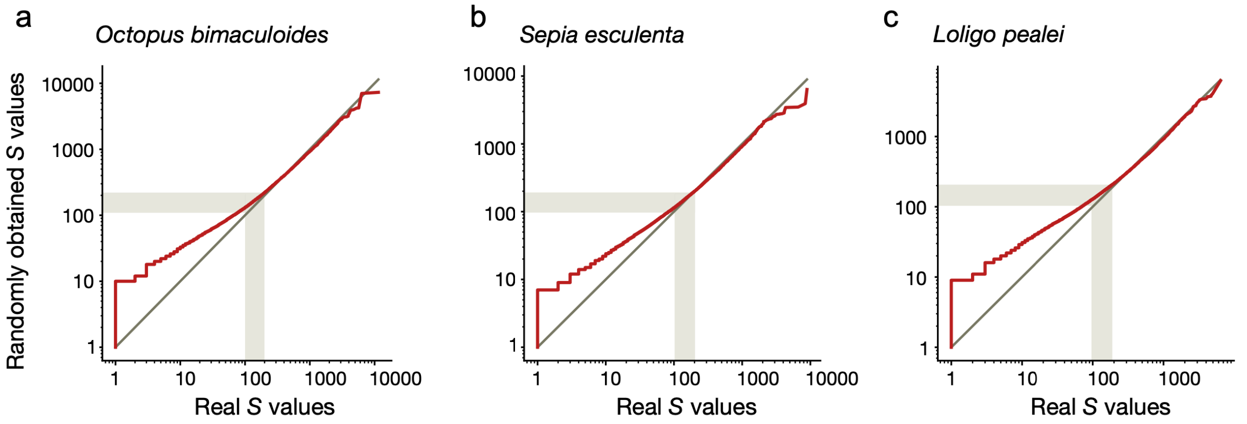


**Supplementary Figure S7 S1 | Clustering of A-to-I editing sites in coleoid transcriptomes.** Red lines show the dependencies between the sorted real and the randomly obtained *S* value sets. The grey lines represent the expected dependence of the form $y=x$. The grey stripes represent the predicted borders of the regions affecting editing sites[1]. **(a)** *O. bimaculoides* **(b)** *S. esculenta* **(c)** *L. pealei*. Notation as in Fig. 6a.

**Supplementary Table S1 | 95% Confidence intervals of differences in base-pairing probabilities between paired editing sites (EE) and three types of control AA-dinucleotides obtained by random sampling.**

| position \ site pair | AA - EE | AE - EE | EA - EE |
|---|---|---|---|
| 1 | 0.0139 to 0.0396 | 0.0573 to 0.0968 | -0.1057 to -0.0399 |
| 2 | -0.0187 to 0.0072 | 0.0494 to 0.0896 | -0.0881 to -0.02 |

#### Supplementary table S2 | SRA identifiers of RNAseq libraries employed in the present study

| SRA ID | Organism | Bioproject |
|---|---|---|
| SRR2045866 | *O. bimaculoides* | PRJNA285380 |
| SRR2045870 | *O. bimaculoides* | PRJNA285380 |
| SRR2047107 | *O. bimaculoides* | PRJNA285380 |
| SRR2047109 | *O. bimaculoides* | PRJNA285380 |
| SRR2047111 | *O. bimaculoides* | PRJNA285380 |
| SRR2047114 | *O. bimaculoides* | PRJNA285380 |
| SRR2047116 | *O. bimaculoides* | PRJNA285380 |
| SRR2047118 | *O. bimaculoides* | PRJNA285380 |
| SRR2047120 | *O. bimaculoides* | PRJNA285380 |
| SRR2047122 | *O. bimaculoides* | PRJNA285380 |
| SRR2048495 | *O. bimaculoides* | PRJNA285380 |
| SRR2048496 | *O. bimaculoides* | PRJNA285380 |
| SRR2048497 | *O. bimaculoides* | PRJNA285380 |
| SRR2048498 | *O. bimaculoides* | PRJNA285380 |

| | | |
|---|---|---|
| SRR2048521 | *O. bimaculoides* | PRJNA285380 |
| SRR2048522 | *O. bimaculoides* | PRJNA285380 |
| SRR2048523 | *O. bimaculoides* | PRJNA285380 |
| SRR2048524 | *O. bimaculoides* | PRJNA285380 |
| SRR2048525 | *O. bimaculoides* | PRJNA285380 |
| SRR2857272 | *O. vulgaris* | PRJNA299756 |
| SRR2857274 | *O. vulgaris* | PRJNA299756 |
| SRR2855904 | *S. esculenta* | PRJNA299756 |
| SRR2856422 | *S. esculenta* | PRJNA299756 |
| SRR1522987 | *L. pealei* | PRJNA255916 |
| SRR1522988 | *L. pealei* | PRJNA255916 |
| SRR1725163 | *L. pealei* | PRJNA255916 |
| SRR1725164 | *L. pealei* | PRJNA255916 |
| SRR1725167 | *L. pealei* | PRJNA255916 |
| SRR1725169 | *L. pealei* | PRJNA255916 |
| SRR1725171 | *L. pealei* | PRJNA255916 |
| SRR1725172 | *L. pealei* | PRJNA255916 |
| SRR1725213 | *L. pealei* | PRJNA255916 |
| SRR1725235 | *L. pealei* | PRJNA255916 |
| SRR1725236 | *L. pealei* | PRJNA255916 |
